# Assessment of Human Renal Transporter Based Drug-Drug Interactions Using Proximal Tubule Kidney-Chip

**DOI:** 10.1101/2022.05.12.491717

**Authors:** Anantha Ram Nookala, Janey Ronxhi, Josiah Sliz, Sauvear Jeanty, Dimitris V. Manatakis, Sushma Jadalannagari, Geraldine Hamilton, Hyoungshin Park, Yu He, Mitchell Lavarias, Gang Luo, Kyung-Jin Jang, Donald Mckenzie

## Abstract

Study of renal transporters is crucial for understanding drug disposition and toxicity, and more importantly, predicting potential drug-drug interactions (DDIs). However, conventional *in vitro* models often fail to predict renal transporter activity and are not scalable to a predictive clinical outcome due to *in vitro-in vivo* discrepancy. Here, we successfully developed a human Proximal Tubule Kidney-Chip model that emulated *in vivo* renal physiology and function to assess renal transporter-based DDIs. Active and improved functionality of key renal transporters including p-glycoprotein (P-gp), multidrug and toxin extrusion (MATE) 1 and 2-K, organic anion transporter (OAT) 1 and 3, and organic cation transporter (OCT) 2 were demonstrated using appropriate probe substrates in Kidney-Chips compared to transwell controls. Moreover, comparative transcriptomic analysis revealed that key efflux and uptake transporters were expressed significantly higher in the Kidney-Chip compared to the transwell. Additionally, key parameters obtained from substrate-inhibitor interactions in the model were used to predict clinical DDIs as well as clearance values, which were closer to *in vivo* clearances. Overall, these results support that the human Proximal Tubule Kidney-Chip can reliably assess the role of human renal transporters in drug disposition and drug interactions, providing a critical tool to assess renal transport *in vitro*.

## Introduction

Renal elimination is an important clearance mechanism for more than 30% of the top 200 prescribed drugs (Morrissey *et al*., 2013). A variety of renal transporters contribute to the renal elimination of several exogenous and endogenous compounds by promoting tubular secretion and tubular reabsorption and thereby, play an important role in their disposition. Transporter activity can be modulated by multiple drugs resulting in the altered disposition, pharmacodynamic response, and development of nephrotoxicity due to drug accumulation (Yin and Wang, 2016). Nephrotoxicity alone accounts for 2% of drug development failures in the pre-clinical stage while rising to 19% in phase 3 clinical trials (Naughton, 2008; Tiong *et al*., 2014). Much of the drug exposure occurs in the proximal tubule as many xenobiotics are secreted by the renal transporters expressed on their apical and basolateral surfaces (Ivanyuk *et al*., 2017). Therefore, evaluation of a new chemical moiety as a victim and perpetrator of renal transporter-mediated drug-drug interactions (DDIs) is recommended by the regulatory agencies around the world, including the U.S. Food and Drug Administration and European Medicines Agency.

Currently, the widely used *in vitro* systems to study the renal transporter-mediated DDIs are human embryonic kidney 293 or Madin-Darby canine kidney (MDCK) cell lines that are transiently transfected with a relevant single transporter (Zhu *et al*., 2012; Müller *et al*., 2018). While these systems are useful to elucidate the role of specific transporter interactions, these are artificial systems overexpressing one specific transporter and cannot be utilized to study the combined effect of more than one transporter. Further, these systems require exhaustive *in vitro* experiments to correlate to *in vivo* interactions (Mathialagan *et al*., 2017). They also lack the expression of several key phenotypic characteristics of renal proximal tubule epithelial cells (RPTECs), which are the prominent cell type within the kidney for drug transport. Within the kidney, RPTECs are exposed to drug-related toxicities as they contain various transporters that accumulate the drugs inside the cells (Filipski *et al*., 2009; Nigam *et al*., 2015; Nieskens and Sjögren, 2019). Therefore, RPTECs are often considered as the gold standard for studying the renal transporter mediated DDIs and nephrotoxicity as they contain the full complement of the renal transporter expression and their gene expression profiles in culture are similar to the *in vivo* renal tissue, unlike the commonly used cell lines. However, RPTEC mRNA transporter expression decreases or is lost upon cryopreservation, and typically, is not restored in conventional 2D static culture models (Van der Hauwaert *et al*., 2014). Different research groups have modified RPTECs to increase their capacity to expand in culture or increase their proliferation at a lower temperature without changing the functional characteristics (Wieser *et al*., 2008; Wilmer *et al*., 2010). Even though these modified RPTECs expressed some of the relevant influx and efflux transporters, they still lacked the expression and functional activity of organic anion transporter 1 (OAT1) and/or OAT3, two of the major drug transporters. Furthermore, RPTECs cultured in a 2D environment lack the expression of appropriate traditional toxicity markers (Li *et al*., 2014), and do not maintain their characteristic polarized membrane structure (Rebelo *et al*., 1992).

The apical surface of the RPTECs is under constant fluid shear stress from the glomerular filtrate which helps with polarization, cytoskeletal rearrangement, expression of apical and basolateral transporters, and localization of tight junction proteins *in vivo* (Duan *et al*., 2008; Ferrell *et al*., 2019). Further, the interstitial space allows for the exchange of various solutes, amino acids, and glucose across the epithelium. Even though the current 2D transwell models allow for the study of bidirectional transport of various substances, it is speculated that the absence of shear stress to be the main reason for the lack of functional cell differentiation leading to the absence of transporter polarization in the current *in vitro* models. These shortcomings with the current *in vitro* models warrant the need for a dynamic system that maintains the RPTECs’ characteristic polarized membrane structure along with the transporter expression/function in culture in order to better model *in vivo* drug transport.

Several research groups have explored the potential of a dynamic microphysiological system that recapitulates the physiology and *in vivo* functions of the proximal tubule (DesRochers *et al*., 2013; Vormann *et al*., 2018). However, limitations occur with these models, including the inability to co-culture multiple cell types or the use of conditionally immortalized cell lines which limit the predictive ability of these models to *in vivo*. In the present study, a dynamic Proximal Tubule Kidney-Chip system was established using human microengineered Organ-on-Chip technology to emulate human physiology and accurately predict renal transporter-mediated DDIs.

## Results

### Mimicking the proximal tubule microenvironment on Kidney-Chip

A Proximal Tubule Kidney-Chip was fabricated to create a micro-engineered environment that recapitulates the proximal tubule section of the kidney. The chip was comprised of two fluidically independent top and bottom channels, separated by a porous polydimethylsiloxane (PDMS) membrane (7 μm pore diameter, ∼3% porosity) coated with ECM (Matrigel and collagen IV) (**Figure 1A**). RPTECs were seeded on the top channel at a density of 2 × 10^6^ cells/mL and renal microvascular endothelial cells (RMVECs) were seeded on the bottom channel at a density of 1 × 10^6^ cells/mL under static conditions. The chips were connected to the pods that contained the RPTEC and RMVEC growth media with a flow rate of 60 μL/hour.

**Figure 1:**
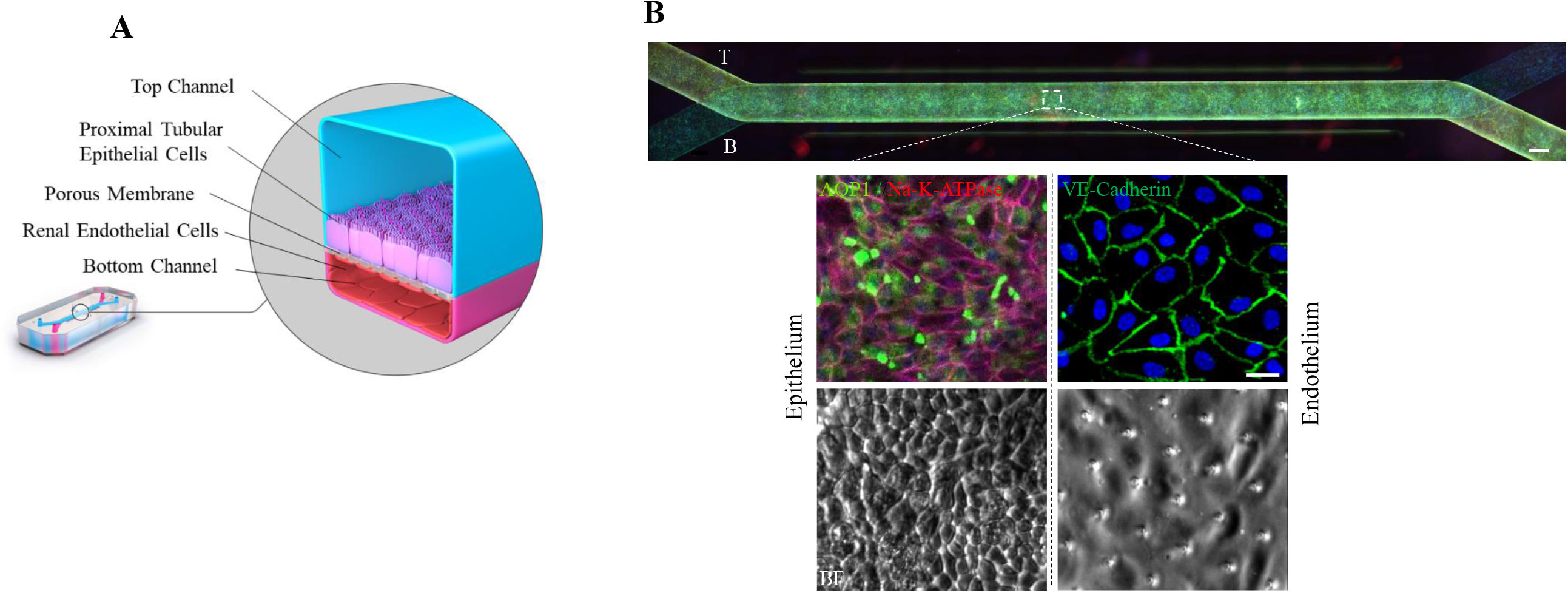
Development and characterization of human Proximal Tubule Kidney-Chip: (A) Cross-section of Kidney-Chip that emulates the structure of the proximal tubule. Proximal tubular epithelial cells are grown in the top channel and renal microvascular endothelial cells are grown in the bottom channel, separated by an extracellular matrix coated porous membrane. (B) Characterization of the proximal tubule epithelial and endothelial cells. Presence of aquaporin-1 and Na^+^/K^+^-ATPase transporters in proximal tubule epithelial cells are indicated by the green fluorescence and red fluorescence, respectively. Confluency and tight junctions of the endothelial cells are indicated by the uniform green staining of VE-cadherin with blue staining nucleus (Hoechst). Bright field (BF) images of epithelium and endothelium on Kidney-Chip after 14 days of culture under flow.

Immunofluorescent staining and imaging were employed to visualize the confluency and polarization by determining the expression of aquaporin 1 (AQP-1), a principal water-transporting protein, and sodium-potassium ATPase (Na^+^/K^+^-ATPase) for the RPTECs and tight junction protein vascular endothelial (VE)-Cadherin for the RMVECs after 14 days in culture. The RPTECs exhibited expression of AQP-1 and uniform staining of Na^+^/K^+^-ATPase marker, indicating that the cells expressed RPTEC markers and formed a confluent monolayer (**Figure 1B**). Similarly, RMVECs showed a uniform and continuous staining of a tight junction protein VE-Cadherin, indicating that the cells formed a uniform and confluent monolayer for 2 weeks in culture (**Figure 1B**). Retained dense monolayer of epithelium and endothelium was also shown by bright field images at 14 days in culture under continuous flow **(Figure 1B**).

### Comparative transcriptomic analysis of transporter gene expression in RPTECs cultured on a Kidney-Chip and transwell

Using RNA-sequencing (RNA-seq), we compared gene expression of the Proximal Tubule Kidney-Chips cultured under constant flow vs conventional static transwells (**Figure 2**). Differential gene expression (DGE) analysis was performed between the Kidney-Chips and transwells which were seeded using the same cell-type composition and subjected to the same experimental conditions for 14 days in culture. Out of the 57,500 genes annotated in the genome, 3,717 were significantly differentially expressed. Among the 3,717 genes in the Kidney-Chip, 1,839 were up-regulated and 1,878 were down-regulated compared to the transwell (**Figure 2B**). Pathway enrichment analysis was performed on the 1,839 up-regulated genes to identify the significantly enriched biological pathways in the Kidney-Chip samples using the Gene Ontology (GO) and Kyoto Encyclopedia of Genes and Genomes (KEGG) database resources. The analyses revealed functional gene sets that significantly clustered under 38 GO and 42 KEGG terms. Apart from kidney-relevant biological processes, these functional gene sets were related to other important biological processes including drug transport, drug metabolic process, fatty acid beta-oxidation, glucose transport, ion transport, oxidation-reduction process (**Figure 2B**). Particularly, efflux transporters including MDR1, MRP2, MRP4, MRP6 and uptake transporters including OAT1, OAT2, OAT3, OAT4, OATP4C1, OCT2, OCTN1, OCTN2, MATE1, and MATE2K were significantly higher expressed in the Kidney-Chip compared to the transwell (**Figure 2C**). Interestingly, efflux transporters which are not expressed in the kidney proximal tubule but instead are localized to the basolateral membranes of the limb of Henle and the distal tubule and collecting duct tubule cells, such as MRP1 and OATP4A1 had decreased expression in the Kidney-Chips compared to transwells (**Figure 2C**) (Masereeuw and Russel, 2012). Similary, the expression levels of the efflux transporter MRP3, which is not expressed in the proximal tubule, was lower in Kidney-Chips compared to transwells but not statistically significant **(Figure 2C)**. These findings demonstrate that the RPTECs in the Kidney-Chip contain higher expression of key drug transporters and other kidney function pathways compared to transwell culture and may be a viable cell culture model to emulate DDIs *in vivo*.

**Figure 2:**
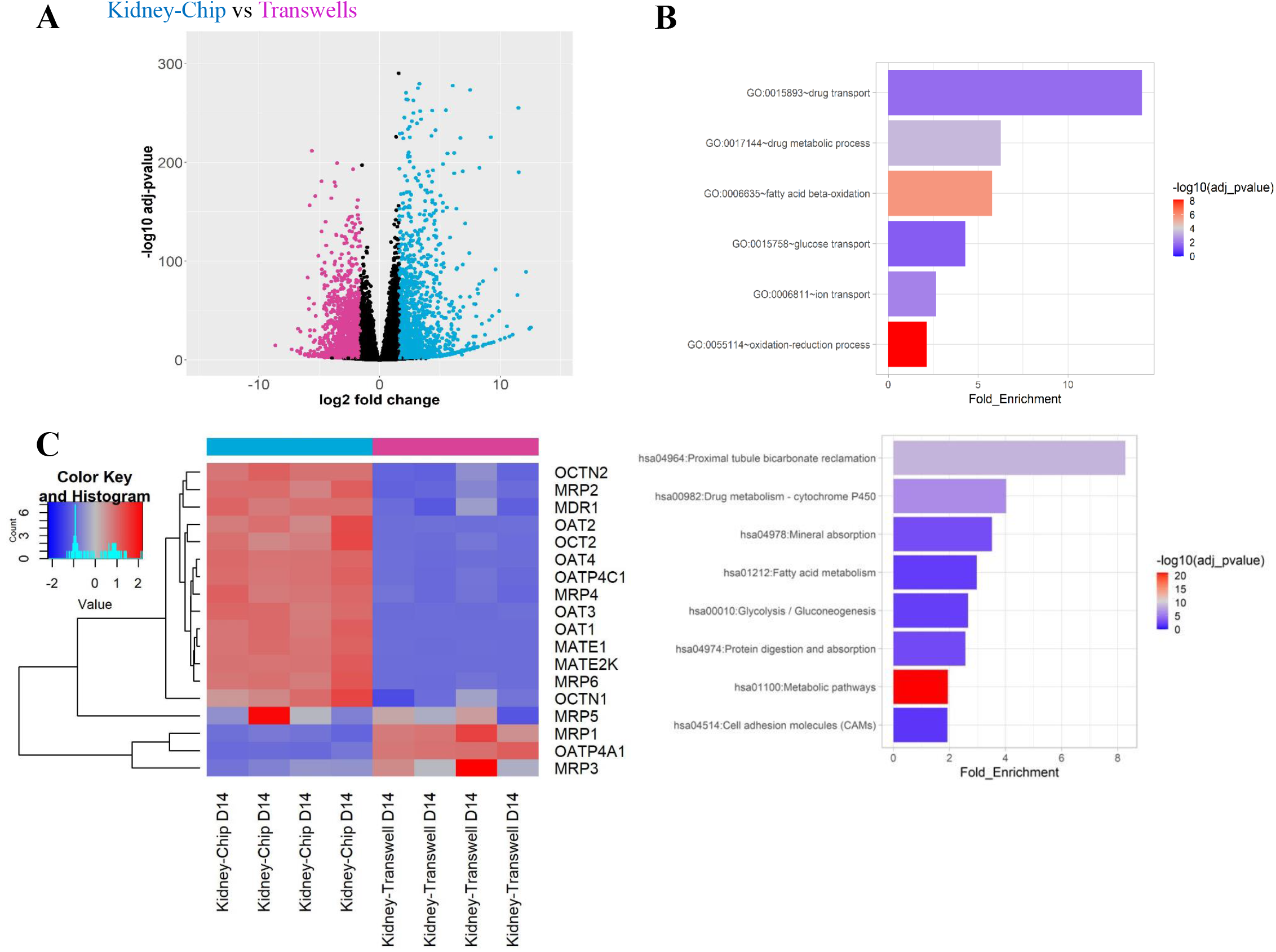
Comparison of Kidney-Chip and transwell transcriptomics data after 14 days in culture: (A) The volcano plot resulted by the DGE analysis between Kidney-Chip and conventional transwell cultures (n=4 samples per group). For the selection of the DE genes we used the following thresholds: adjusted p-value< 0.01 and |Log2 (FoldChange)| > 1.6. The identified up- and down-regulated genes are highlighted in cyan and magenta color, respectively. (B) List of biological processes identified by KEGG pathway and Gene Ontology (GO) enrichment analysis using the up-regulated genes resulted by the differentially gene expression analysis between Kidney-Chip and conventional transwell cultures. (C) Heatmap was generated to examine particular genes of efflux transporters including MDR1, MRP1, MRP2, MRP3, MRP4, MRP6 and uptake transporters including OAT1, OAT2, OAT3, OAT4, OATP4A1, OATP4C1, OCT2, OCTN1, OCTN2, MATE1, and MATE2K between Kidney-Chip and transwell cultures.

### Comparison of transporter function in Proximal Tubule Kidney-Chip and transwell co-culture

The functional activity of various transporters on Day 14 post-seeding on Kidney-Chip and transwell was compared and shown in **Figure 3B** and C. Specific probe substrates used to evaluate the functional activity of transporters included ^3^H-digoxin for P-gp, ^14^C-TEA and ^14^C-metformin for OCT2, MATE1, and MATE-2k activity, and ^14^C-para-aminohippuric acid (PAH) and ^14^C-adefovir for OAT1. The location of the transporters in the proximal tubule along with the direction in which the substrates are transported is presented in **Figure 3A**. Chips were dosed in either top channel/apical side (A) or bottom channel/basal side (B) under flow (Error! Reference source not found.).

**Figure 3:**
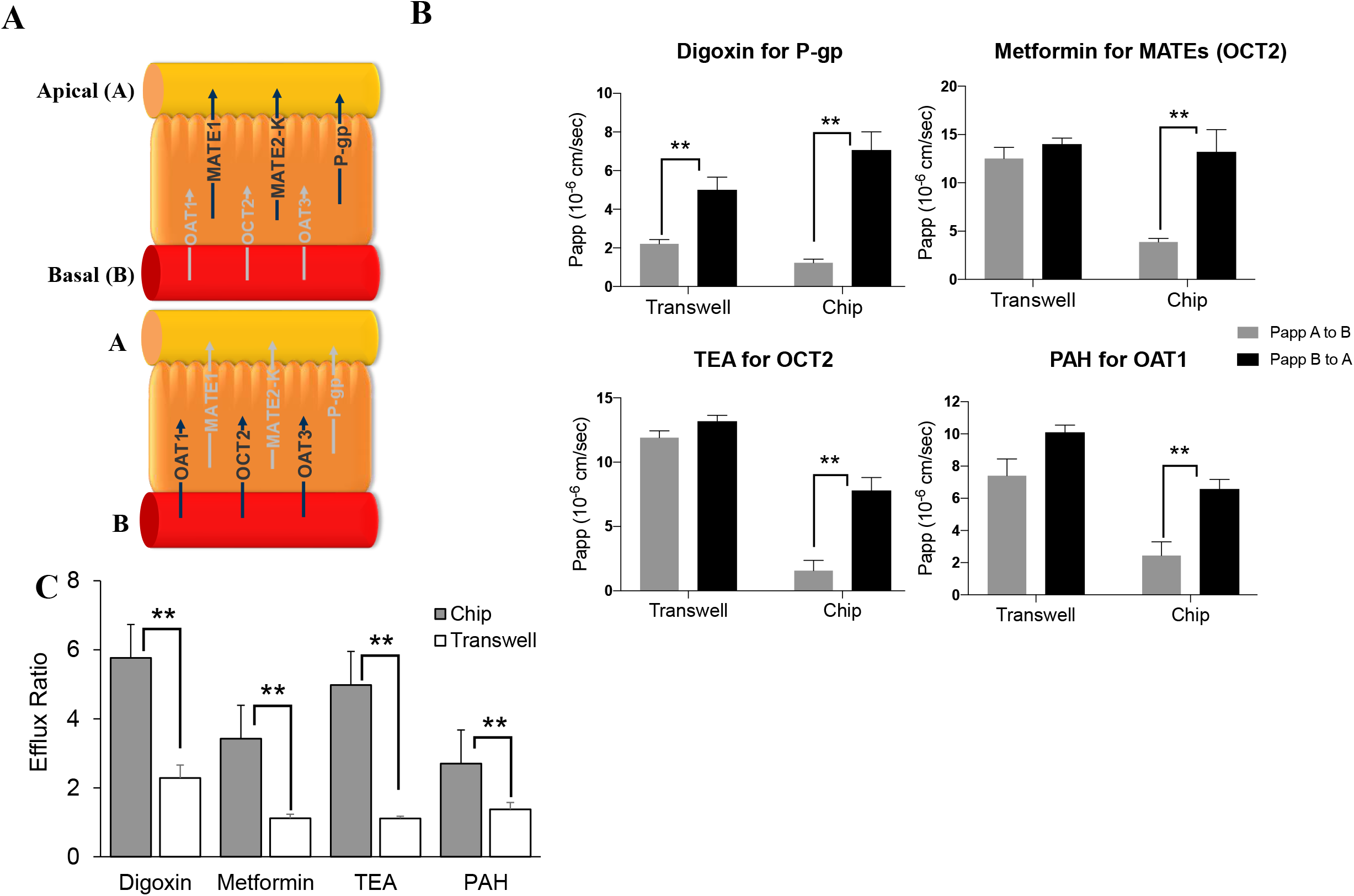
Comparison of the transporter activity on Kidney-Chip versus transwell after 14 days in culture: (A) Schematic of the location and transport direction of apical (MATE1, MATE-2K, and P-gp) and basolateral transporters (OAT1, OAT3, and OCT2). (B) Apparent permeabilities (P_app_) for various probe substrates, including digoxin (P-gp), metformin (MATEs and OCT2), tetraethyl ammonium (OCT2), and p-amino hippuric acid (OAT1) are presented. Apparent permeability from A to B and B to A directions are represented in gray and black solid bars, respectively. (C) Efflux ratios of the probe substrates are presented for Kidney-Chip and transwell by gray and white solid bars, respectively (n = 3 independent chips). Statistical analyses were performed by Student’s *t*-test and ** denotes *P*-value of ≤ 0.01.

#### Digoxin

The apparent permeability (P_app_) of ^3^H-digoxin from A to B and B to A was ≥2.21 × 10^−6^ and ≥5.00 × 10^−6^ cm/sec, respectively in transwell; ≥1.23 × 10^−6^ and ≥7.06 × 10^−6^ cm/sec, respectively in Kidney-Chips. The efflux ratio of ^3^H-digoxin was 5.76 in Kidney-Chip, approximately 2.5-fold of that in transwell (2.28).

#### Metformin

The P_app_ of ^14^C-metformin from A to B and B to A was ≥12.50 × 10^−6^ and ≥14.00 × 10^−6^ cm/sec, respectively in transwell; ≥3.87 × 10^−6^ and ≥13.20 × 10^−6^, respectively in Kidney-Chips. The efflux ratio of ^14^C-metformin was 3.42 in Kidney-Chip, approximately 3-fold of that in transwell (1.12).

#### TEA

The P_app_ of ^14^C-TEA from A to B and B to A was ≥11.90 × 10^−6^ and ≥13.20 × 10^−6^ cm/sec, respectively in transwell; ≥1.57 × 10^−6^ and ≥7.80 × 10^−6^, respectively in Kidney-Chips. The efflux ratio of ^14^C-TEA was 4.98 in Kidney-Chip, approximately 4.5-fold of that in transwell (1.11).

#### PAH

The P_app_ of ^14^C-PAH from A to B and B to A was ≥7.40 × 10^−6^ and ≥10.10 × 10^−6^ cm/sec, respectively in transwell; ≥2.44 × 10^−6^ and ≥6.58 × 10^−6^, respectively in Kidney-Chips. The efflux ratio of ^14^C-PAH was 2.7 in Kidney-Chip, approximately 2.4-fold of that in transwell (1.13).

A similar trend was observed in Day 8 in culture for both transwell and chip groups, with slight increases of efflux ratio over time in the Kidney-Chip for all substrates (**Supplementary Figure 3**). These results confirmed the transcriptomics data. These results also indicated that the key renal transporters P-gp, OCT2, MATE1, MATE2-k, and OAT1 were functionally active between Day 8 and Day 14 in Kidney-Chip when the RPTECs and RMVECs were co-cultured on the Kidney-Chip, whereas only P-gp was active to some extent and rest renal transporters were inactive in the transwell. Altogether the data suggest that the Proximal Tubule Kidney-Chip performs key functions of *in vivo* kidney drug transport.

### Prediction of clearance from Kidney-Chip and Transwell efflux ratio

To better understand if the Proximal Tubule Kidney-Chip could predict drug transport *in vivo*, transporter data obtained in the absence of inhibitors including efflux ratios, predicted clearance, *in vivo* clearance cited from literature reports, and contribution of distal tubule reabsorption to kidney clearance are summarized in **Table 1**. Renal clearance obtained clinically or predicted from Kidney-Chip and Transwell are presented graphically in **Figure 4**.

**Table 1:** Summary of apparent permeability, efflux ratio, and clearance values calculated using Kidney-Chip and transwell.

| Substrate | Transwell Efflux Ratio | Kidney-Chip Efflux Ratio | Total in vivo Clearance (mL/min) | Transwell Predicted Clearance |  | Kidney-Chip Predicted Clearance |  | Distal Tubule Reabsorption |
| --- | --- | --- | --- | --- | --- | --- | --- | --- |
|  |  |  |  | mL/min | % of in vivo | mL/min | % of in vivo |  |
| Adefovir | - | 2.01+/-0.11 | 259 ± 0.61 | - |  | 221 ± 13 | 85 | Minimal |
| Digoxin | 2.28+/-0.38 | 8.34+/-0.83 | 119 | 195 ± 33 | 164 | 516 ± 70 | 434 | Significant |
| Metformin | 1.12+/-0.12 | 3.26+/-0.41 | 533 ± 21 | 134 ± 14 | 25 | 341 ± 47 | 64 | Minimal |
| PAH | 1.37+/-0.20 | 4.84+/-0.57 | 540 | 135 ± 20 | 25 | 390 ± 54 | 72 | Minimal |
| TEA | 1.11+/-0.06 | 6.95+/-0.60 | Unavailable | 133 ± 7.2 |  | 568 ± 68 |  | - |

**Figure 4:**
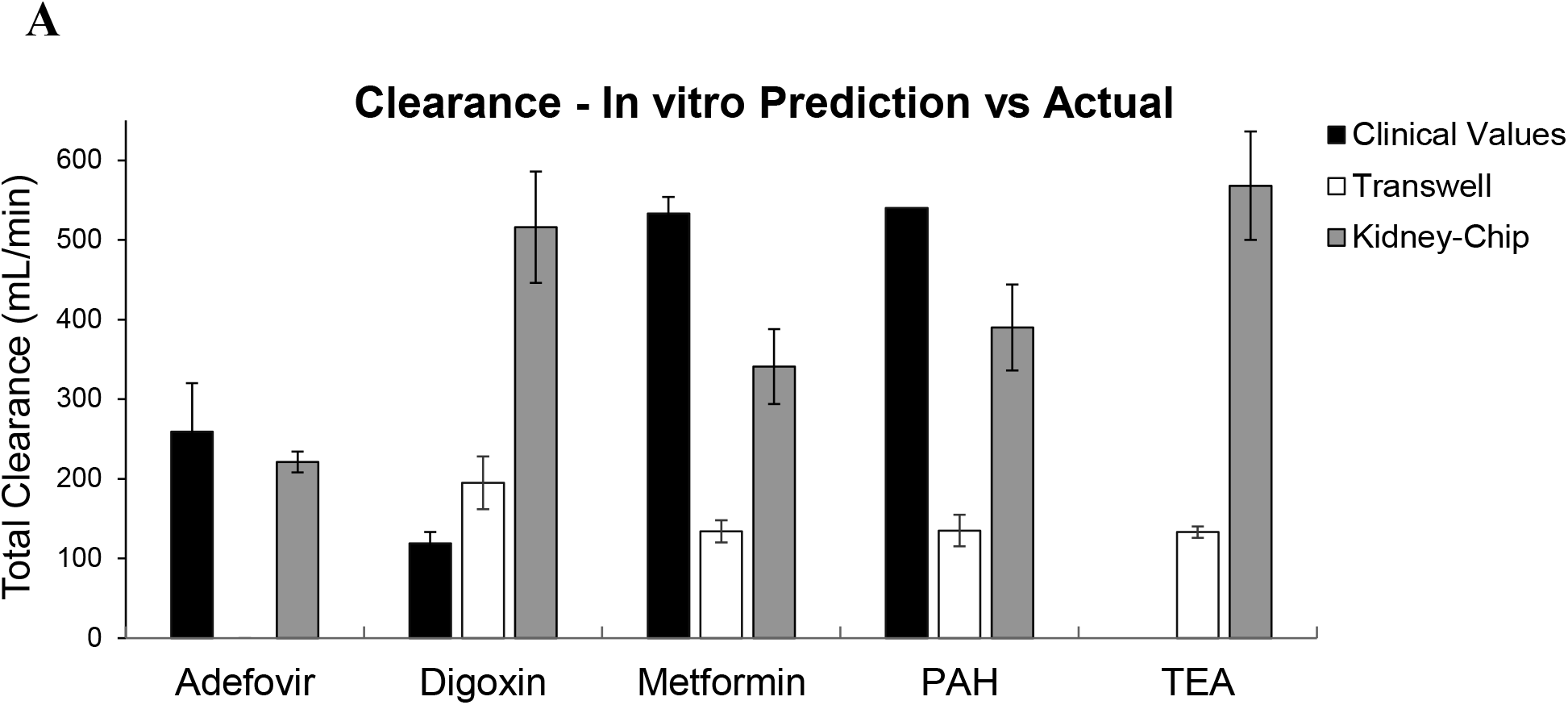
Comparison of *in vitro* predicted clearance values to *in vivo*: Clearance (mL/min) was calculated for adefovir, digoxin, metformin, p-amino hippuric acid, and tetra ethyl ammonium using data obtained from Kidney-Chip and transwell and was compared against the *in vivo* values observed in human subjects. Kidney-Chip, transwell, and human *in vivo* clearance values are represented by gray, white, and black solid bars, respectively.

Compared with the *in vivo* clearance, the clearance predicted from transwell was 164% for digoxin, 25% for metformin, and 25% for PAH, whereas the clearance predicted from Kidney-Chip was 85% for adefovir, 434% for digoxin, 64% for metformin, and 72% for PAH. Much higher *in vitro* clearance for digoxin in Kidney-Chip was likely due to the model lacking distal tubule reabsorption which markedly reduces *in vivo* clearance. Slightly lower *in vitro* clearances for adefovir, metformin, and PAH might be due to lower transporter activities in the *in vitro* models or due to the interindividual differences in the transporter activities. Significantly, the clearances predicted from Kidney-Chip for adefovir, metformin, and PAH were closer to *in vivo* clearances and much better than those predicted from transwell.

### *In vitro* Renal Transporter Mediated DDI

As various drugs are shown to affect the secretion of certain xenobiotics across the RPTECs by inhibiting the transporter function resulting in drug-drug interaction we wanted to test if the Kidney-Chip model would show similar interactions. In the Kidney-Chip model, the tested transporters were shown to be functionally active between Day 8 and Day 14; accordingly, various renal transporter mediated DDI experiments were performed between Days 8 and 14.

#### Digoxin-Quinidine DDI

Quinidine (1, 5, and 25 µM) inhibited P-gp-mediated transport of digoxin (1 µM) in a concentration-dependent manner (**Figure 5A**). The efflux ratio was approximately 1 at 25 µM quinidine. An IC_50_ value was estimated to be 0.723 µM.

**Figure 5:**
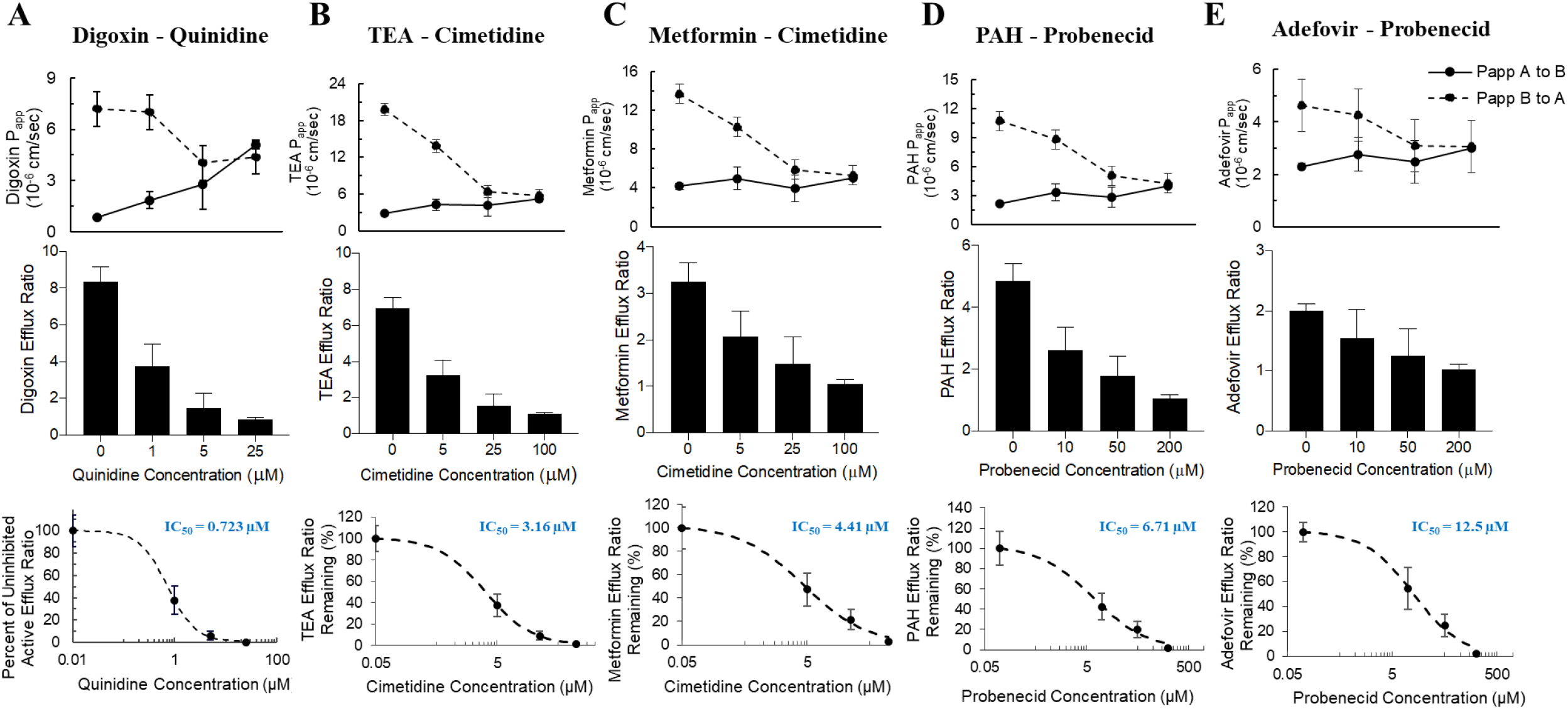
Utilizing Kidney-Chip to study various renal transporter mediated drug-drug interactions: RPTECs and RMVECs were cultured on Kidney-Chip for 8 to 14 days under a flow rate of 60 µL/hr. Various concentrations of solvent control or inhibitor solutions were added to the top and bottom channels 1 hour prior to the addition of the probe substrates and incubated for 3 hours. Different substrate-inhibitor combinations included were (A) Digoxin-quinidine, (B) TEA-cimetidine, (C) Metformin-cimetidine, (D) PAH-probenecid, and (E) Adefovir-probenecid. Apparent permeability (P_app_) from A to B and B to A directions are represented by solid line and dashed line, respectively. Efflux ratios of each substrate at different inhibitor concentrations are presented in a bar graph. Percent of uninhibited active efflux ratio against inhibitor concentrations was plotted and the corresponding IC_50_ values were calculated (n = 3 independent chips).

#### TEA-Cimetidine DDI

Cimetidine (5, 25, and 100 µM) inhibited OCT2- and MATEs-mediated transport of TEA (1 µM) in a concentration-dependent manner (**Figure 5B**). The efflux ratio was approximately 1 at 100 µM cimetidine. The IC_50_ value was estimated to be 3.16 µM.

#### Metformin-Cimetidine DDI

Cimetidine (5, 25, and 100 µM) also inhibited OCT2- and MATEs-mediated transport of metformin (1 µM) in a concentration-dependent manner (**Figure 5C**). The efflux ratio was approximately 1 at 100 µM cimetidine. The IC_50_ value was estimated to be 4.41 µM.

#### PAH-Probenecid DDI

Probenecid (10, 50, and 200 µM) inhibited OAT1-mediated transport of PAH (1 µM) in a concentration-dependent manner (**Figure 5D**). The efflux ratio was approximately 1 at 200 µM probenecid. The IC_50_ value was estimated to be 6.71 µM.

#### Adefovir-Probenecid DDI

Probenecid (10, 50, and 200 µM) inhibited OAT1-mediated transport of adefovir (1 µM) in a concentration-dependent manner **(Figure 5E**). The efflux ratio was approximately 1 at 200 µM probenecid. The IC_50_ value was estimated to be 12.5 µM.

All the IC_50_ values estimated above can be taken to approximate the inhibition constant (K_i_) as all the probe substrates were used well below its transport Michaelis-Menten Constant (k_m_). These results indicated that DDIs mediated by key renal transporters can be studied using this Proximal Tubule Kidney-Chip model.

### Simulation of Clinical DDI Studies

Clinical pharmacokinetics of digoxin, metformin, and adefovir were simulated using a one-compartment clearance model. The simulation was performed for each drug in the absence and presence of respective inhibitors (**Figure 6A, B, and C**). Subsequently, clinical impact (CI) was calculated for each DDI (**Table 2**). Predicted CI was also calculated based on the results obtained from Kidney-Chip model (**Table 2**). Inhibition studies in transwell were not performed and no comparison to *in vivo* was attempted since the transwell data was not suitable for this analysis. Indeed, because the efflux ratios were low, no significant difference was expected between the predicted uninhibited and completely inhibited case, but rather both predictions would be primarily dominated by glomerular filtration.

**Figure 6:**
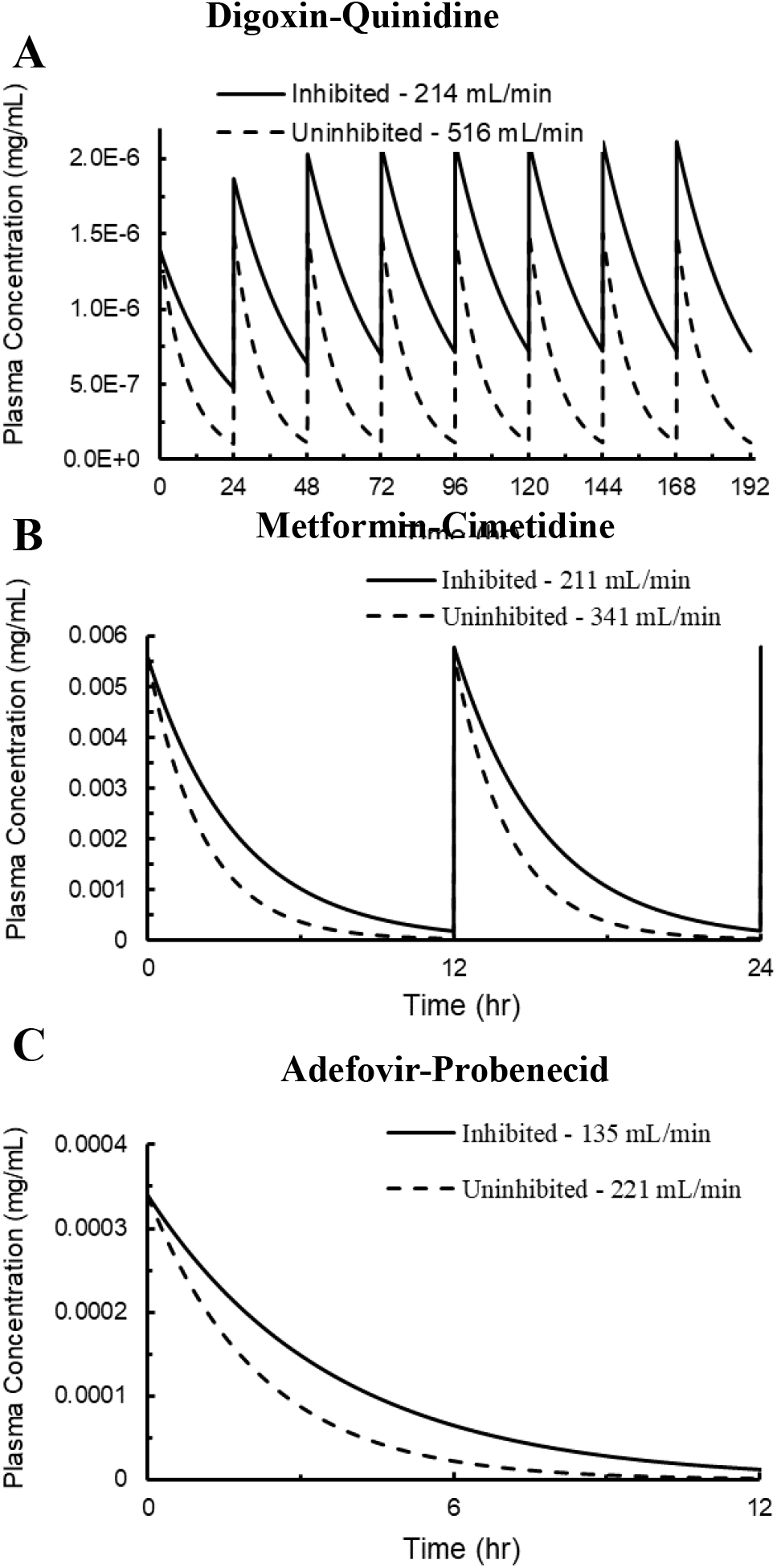
Simulation of clinical drug-drug interactions using Kidney-Chip: Simulation of plasma concentration-time profiles of (A) Digoxin-quinidine, (B) Metformin-cimetidine, and (C) Adefovir-probenecid. Solid lines and dashed lines represent clearance in the absence and presence of inhibitor, respectively.

**Table 2:** Summary of Kidney-Chip predicted clinical impact to actual clinical impact on AUC and Cmax.

| Implicated<br>Transporter<br>Family | Perpetrator | Victim | Clinical<br>Impact | Predicted<br>Clinical Impact |
| --- | --- | --- | --- | --- |
| P-gp | Quinidine | Digoxin | Cl<sub>r</sub> ↓54%<br>AUC ↑166% | Cl<sub>r</sub> ↓59%<br>AUC ↑128% |
| OCT/MATE | Cimetidine | Metformin | Cl<sub>r</sub> ↓27%<br>AUC ↑50% | Cl<sub>r</sub> ↓20-38%<br>AUC ↑26-60% |
| OAT | Probenecid | Adefovir | Not Reported<br>AUC ↑82% | Cl<sub>r</sub> ↓39%<br>AUC ↑64% |

Importantly, while model-generated concentration profiles are shown with time for metformin adefovir, and digoxin, the absolute values may be skewed due to either under-prediction or over-prediction of kidney clearance (see **Table 1** for level of over/underprediction). For example, digoxin clearance is over-predicted from Kidney-Chip efflux ratios, likely due to the contribution of distal tubule reabsorption in minimizing total excretion *in vivo*. While this does result in an over-prediction of clearance, it does not result in skewing of predicted *shifts* in clearance due to inhibition, since distal tubule reabsorption is dependent on the fraction reabsorbed relative to the summation of glomerular clearance and proximal tubule secretion clearance. Thus, so long as estimates of glomerular and proximal tubule secretion are correct, inhibition of proximal tubule secretion will result in the same shift in clearance and area under the C_p_ curve regardless of whether distal tubule reabsorption is correctly estimated.

#### Digoxin-Quinidine DDI (Figure 6A)

We modeled a clinical DDI study examining the effects of quinidine inhibition of digoxin active transport on the clearance of digoxin from systemic blood circulation. The study consisted of an 8-day oral dosing regimen, with digoxin being administered either alone, once daily for 8 days at 0.4 mg/dose or co-administration with quinidine twice daily at 600 mg/dose for 8 days, in a random crossover study design (Rameis, 1985). Clinical results were reported for the observed shift in total clearance and shift in Area Under the Curve (AUC) for a population of 6 healthy volunteers. The clearance model predicted a decrease in total kidney clearance of 59% due to quinidine inhibition, with the model run using the C_ss_ of quinidine. This compared well to the clinically observed decrease in total clearance of 54%. AUC was predicted to increase by a very appreciable 128%, which was in line with observed clinical values, where AUC increased by an average of 166%. As can be seen in Figure 6A, with multiple doses, circulating digoxin concentration increases further from the uninhibited case as time progresses and additional doses are administered, which further increases the exposure or AUC.

An additional four clinical DDI studies were modeled for digoxin-quinidine DDI, each following a similar experimental design (results not shown in Figure 6) (Hager *et al*., 1981; Ochs *et al*., 1981; Fenster *et al*., 1984). The studies each consisted of an initial single 1 mg IV dose of digoxin to assess uninhibited clearance in the subjects, followed by between 4 and 11 days of exposure to quinidine via 4X daily oral administration of 200 mg tablets, followed by another single 1mg IV dose of digoxin. Clinical results were only reported for the observed shift in total clearance for a population of between 6 and 7 healthy volunteers, depending on the study. The clearance model predicted a decrease in total kidney clearance of 46 % due to quinidine inhibition, with the model run using the C_ss_ of quinidine. This compared well to the average clinically observed decrease in total clearance of also 46 %, indicating very good agreement between the model and clinical results.

#### Metformin-Cimetidine DDI (Figure 6B)

We modeled a clinical DDI study examining the effects of cimetidine inhibition of metformin active transport on the clearance of metformin from systemic blood circulation (Somogyi *et al*., 1987; Wang *et al*., 2008). The study consisted of a 10-day oral dosing regimen, with metformin being administered alone, twice daily for 5 days at 250 mg/dose, followed by co-administration of metformin with cimetidine for another 5 days, which was also administered twice daily at 400 mg/dose. Clinical results were reported for the observed shift in total clearance and shift in AUC for a population of 7 healthy volunteers. The clearance model predicted a decrease in total kidney clearance of between 20 – 38% due to cimetidine inhibition, depending on whether the model was run using cimetidine concentration at steady-state (C_ss_) or peak circulating concentration (C_max_). This compared well to the clinically observed decrease in total clearance of 27%. Similarly, AUC was predicted to increase by between 26 – 60% and was observed to increase in the clinical setting by an average of 50%, indicating good agreement between the model and clinical results.

#### Adefovir-Probenecid DDI (Figure 6C)

We also modeled a clinical DDI study examining the effects of probenecid inhibition of adefovir active transport on the clearance of adefovir from systemic blood circulation (Maeda *et al*., 2014). The study consisted of a single day oral co-dose of probenecid at 10mg and adefovir at 1500 mg. Clinical results were reported for the observed shift in total clearance and shift in AUC for a population of 6 healthy volunteers. The clearance model predicted a decrease in total kidney clearance of 39% due to probenecid inhibition, with the model run using the peak circulating probenecid concentration (C_max_), as a concentration at steady-state cannot be computed for single dosage regimen. The shift in clearance in the clinical setting was not reported for this study. AUC was predicted to increase by 64% and was observed to increase in the clinical setting by an average of 82%, indicating good agreement between the model and clinical results.

Overall, we observed a good agreement with simulations obtained using the parameters from the Kidney-Chip compared to the literature values for digoxin, metformin, and adefovir. These results also indicated that the one compartmental clearance model was adequate for predicting the clinical pharmacokinetics (clearance and AUC) of digoxin, metformin, and adefovir in the presence and absence of inhibitors.

## Discussion

The kidney is an organized tissue comprised of different cell types surrounded by intricate capillary networks and extracellular matrix. Various cell lines routinely used to study the transporter mediated drug interactions and drug induced nephrotoxicity are of non-human origin and are utilized in conventional 2D models without fluid flow (Wilmer *et al*., 2016). Given the limitations of culturing primary epithelial cells on a 2D model, there is a need to develop a kidney microphysiological systems (MPS) that recapitulates the dynamic environment, structure and function of kidney *in vivo*. Previously, several groups have developed bioengineered 3D kidney tissue models to study renal transport, nephrotoxicity, and drug interactions (Humes *et al*., 1999; DesRochers *et al*., 2013; Vormann *et al*., 2018). However, the use of immortalized cell lines or inability to co-culture is a limitation of these models. Our model overcomes these limitations to successfully co-culture primary RPTECs and RMVECs exposed to unidirectional flow on both sides emulating the dynamic microenvironment of proximal tubule of the kidney in the microfluidic chip system.

Fluid shear stress regulates the expression of tight junction proteins on epithelial cells which restrict the leakage of solutes, amino acids, glucose, and nutrients (Maggiorani *et al*., 2015). The RPTECs and RMVECs of Kidney-Chip cultured under fluid shear stress retained their characteristic morphology with the localization of tight junction proteins for the entire culture duration (Duan *et al*., 2008). Fluid shear stress induced phenotypic polarization of the epithelial cells was confirmed by staining for Na^+^/K^+^-ATPase and P-gp on the basolateral and apical surfaces, respectively. In contrast, the cells cultured on a conventional 2D transwell condition do not retain the characteristic cuboidal morphology and have leaky junctions (Hoppensack *et al*., 2014). Numerous cilia extending into the lumen on the surface of the RPTECs play important roles in calcium signaling, endocytosis, and mechanosensing (Pazour and Witman, 2003; Raghavan *et al*., 2014). We previously used immunocytochemistry and scanning electron microscopy analysis to demonstrate the expression of primary cilia on the surface of RPTECs (Jang *et al*., 2013). One of the important functions of the proximal tubule is reabsorption of serum albumin secreted into the glomeruli through megalin- and cubulin-mediated endocytosis (Merlot *et al*., 2014) (Zhai *et al*., 2000; Nielsen *et al*., 2016). We also previously reported that RPTECs cultured on the Kidney-Chip under fluid shear stress retained the apical endocytic capacity by quantifying the uptake of FITC-labeled albumin (Jang *et al*., 2013).

RPTECs mediate the transfer of endogenous and exogenous compounds from lumen to blood or blood to lumen through functionally distinct apical and basolateral membrane transporters (International Transporter Consortium *et al*., 2010). To date, there are only few *in vitro* models that recapitulate the kidney structure and function *in vitro* (Maggiorani *et al*., 2015; King *et al*., 2017). However, none of the models have completely characterized the expression of renal drug transporters and their potential to assess drug-drug interactions. Previously, research groups have indicated that fluid shear stress can increase the expression of transporters in mouse proximal tubule cells (Wang *et al*., 2017) or MDCK cell line cultured in microfluidic biochip for up to 96 hours (Snouber *et al*., 2012). Our transcriptomic analysis showed that fluid shear stress induced significant changes in the gene expressions of various transporters and enzymes involved in various physiologic and metabolic functions. Compared to the RPTECs and RMVECs cultured on a conventional transwell, we observed a significant upregulation in the expression of several key transporters, including P-gp, OAT1/3, MRP2/4, OCT2, and OCT2 when cultured on the Kidney-Chip.

Given the transcriptome analysis, we compared the functional activity of various transporters in Kidney-Chip to the transwell control. Except for P-gp, the functional activity of various transporters was absent in RPTECs cultured on the transwell plate. The flow retained the functional activity of various transporters in the Kidney-Chip as demonstrated by the uptake and directed efflux of several probe substrates. We also confirmed dose-dependent drug interactions with various transporters using the regulatory agency recommended probe substrates and inhibitors. There has been some limited research regarding the effect of fluid flow on the functionality of renal transporters; P-gp and MRP2/4 in immortalized PTECs (Vriend *et al*., 2020), OAT1/3 in human PTECs (Weber *et al*., 2016), OAT in porcine PTECs (Humes *et al*., 1999), and OCT2/MATE1 in transfected MDCK cells (Jayagopal *et al*., 2019). However, none of these models have adequately characterized the functional activity of all the relevant human renal transporters within a single model and have not utilized the co-culture of epithelial and endothelial cells which is a closer representation of kidney tubular environment *in vivo*. Further, some of these models used cells from non-human origin or immortalized cells that makes the extrapolation to human in vivo data difficult.

As a demonstration of the Kidney-Chip’s utility to predict clinically relevant DDI outcomes, we used the results from the Kidney-Chip dose-dependent DDI studies to model the impact of *in vivo* administration of prototypic inhibitors on the rate of elimination and total exposure to co-administered substrates of proximal tubule active transport. When following the same dosing regimen as outlined in clinical studies, we were able to accurately predict shifts in clearance and AUC for the three victim/perpetrator combinations for which clinical data was available from the University of Washington Drug Interaction Database. These clinically relevant parameters were confirmed for metformin-cimetidine and adefovir-probenecid in a repeat-dose and single-dose regimen, respectively, and for digoxin-quinidine in both a single-dose and repeat-dose regimen study design. Altogether, our data suggests that the Kidney-Chip model more accurately predicts *in vivo* drug clearance than other *in vitro* kidney models.

Future work will, in part, focus on adding additional complexity to the computational models used to support extrapolation of Kidney-Chip data to *in vivo* parameters/clinical outcomes to better predict *in-vivo* results. Specifically, the one-compartment clearance model could be expanded to a multi-compartment physiologically-based pharmacokinetic model to capture important compound kinetics such as elimination of compound from other clearance pathways such as liver metabolism. Incorporation of these other clearance mechanisms would improve the direct applicability of model estimations of clearance and, consequently, the direct applicability to decisions in the clinical setting without needing to consider non-renal clearance mechanisms separately. For the compounds evaluated here, kidney clearance was the major route of compound elimination from the body and incorporation of liver clearance, for example, was not necessary (Goodman *et al*., 2011).

Additional complexity could also be beneficial to the approach used to extrapolate *in vitro* efflux ratios to *in vivo* clearances (Scotcher *et al*., 2016). For example, with the current extrapolation procedure, the contribution of distal tubule reabsorption is not considered quantitatively, instead only indicated whether the compound is known to reabsorb appreciably. Further modeling is required to appropriately capture the time-dependent nature of reabsorption, incorporate the differences in kidney physiology seen in the population, including differences in diet, which can cause a range in pH from 4.5 – 8.0 within the tubule and contribute to the extent of the equilibrium condition achieved before filtrate enters the collecting ducts. Tubule pH can dramatically affect ionization of compound within the tubule and, therefore, the driving-force or extent of reabsorption (Scotcher *et al*., 2016; Mathialagan *et al*., 2017). For the compounds examined in these studies, only digoxin is known to be extensively reabsorbed in the distal tubule and thus only digoxin clearance was over-estimated with this approach. Albeit, even in the case of digoxin, decreases in clearance and increases in AUC due to inhibition could be well predicted since shifts in clearance (in terms of percent change) are not impacted by the absolute value of fraction reabsorbed incorporated in the model. Nonetheless, future work is needed to create a more complete physiologically-based model of the proximal tubule and distal tubule, which incorporates the development of gradients along the length of tubule and will improve the power of the model by enabling more accurate and robust prediction of *in vivo* clearance, as opposed to the current approach which only assesses upper and lower limits.

Despite the current use of classic computational models, Kidney-Chip modeling is able to give a better quantitative prediction of *in-vivo* DDIs in the clinical setting and an important improvement over the estimation of uninhibited clearance in comparison to transwell. Expected shifts in clearance and AUC due to DDIs in the clinical setting were predicted well by leveraging Kidney-Chip victim-perpetrator dose-response data and simple/classical computational models. Information such as this would be of extreme value for quantitatively assessing the impact of DDIs on novel pharmaceutical exposure. Shifts in exposure can have important clinical outcomes, especially in the case of multi-dose regimen, where a significant increase in the AUC is possible and the risk of chronic toxicity amplified. The Kidney-Chip appears well-positioned to fill the niche of quantifying the severity of DDI risk and defining concentrations of inhibitor and particular dosing regimen where these DDIs will be important.

In summary, we have developed and shown for the first time a microphysiological system that emulates the proximal tubule portion of the kidney to study renal transporter mediated DDIs. The clinical pharmacokinetic parameters obtained from the Kidney-Chip were close to the clinical outcomes, markedly improving prediction of *in vivo* outcomes compared to the current standard of transwells.

## Materials and methods

### Materials

^14^C-Adefovir, ^3^H-digoxin, ^14^C-metformin,, ^14^C-para-aminohippuric acid, and ^14^C-tetraethyl ammonium were purchased from American radiolabeled chemicals (St. Louis, MO). Cimetidine, probenecid, quinidine, and collagen IV were purchased from Millipore Sigma (St. Louis, MO). Heat inactivated fetal bovine serum (HI FBS) and trypan blue were procured from Gibco Life Technologies (Waltham, MA). Cryopreserved primary human RPTECs (donor lot CC-2553, female, 44 years old), REBM renal epithelial cell growth basal medium, REGM single quot kit, and amphotericinB/gentamycin were procured from Lonza (Basel, Switzerland). Cryopreserved primary RMVECs (donor lot 128.02.01.02.0R, pooled), CSC basal medium, and culture boost were procured from cell systems (Kirkland, WA). Matrigel, DPBS, and transwell plates were purchased from corning (Corning, NY).

### Cell culture

RMVECs were cultured in CSC basal medium supplemented with culture boost, HI FBS, and amphotericin B/gentamycin. RPTECs were cultured in REBM renal epithelial cell growth basal medium supplemented with human epidermal growth factor, insulin, hydrocortisone, transferrin, human triiodothyronine, human epinephrine, HI FBS, and amphotericin B/gentamycin. Cells were maintained in an incubator at 37°C, 5% CO_2_ and saturated humidity for 4 days before plating into extracellular matrix (ECM) coated chips or transwell plates.

The chips (S-1 Chips, Emulate Inc.) were activated with a proprietary ER-1 solution (Emulate Inc.) under UV light for 15 minutes. The chips were incubated overnight at 37°C with ECM mixture containing collagen IV (50 µg/mL) and matrigel (100 µg/mL). RMVECs were plated into the bottom channel at a density of 2 × 10^6^ cells/mL and the chips were inverted for 3 hours to facilitate attachment to the porous membrane that separates the two parallel channels in the chip. The chips were flipped back to the upright position, followed by the addition of RPTECs into the top channel at a density of 1 × 10^6^ cells/mL A gentle gravity wash with the fresh media was performed after the cells have fully attached (approximately 3 hours post-seeding) to ensure that nutrients were replenished. Following day, the cells were appropriately washed with the medium and were connected to the Pods and Human Emulation System (Emulate Inc.). The flow rate was set to 60 µL/hour for the top and bottom channels. The medium was appropriately replenished for the duration of the experiment.

A parallel experiment using co-cultured transwell control (polyester membrane, pore size 0.4 µm) was conducted to investigate flow effect on RPTEC function in the Kidney-Chip. RMVECs were suspended in CSC media at a density of 0.5 × 10^6^ cells/mL. The transwell insert was inverted and 100 µL of the cell suspension was added to the surface of each transwell insert. After 3 hours, the transwell was reverted to its original position. RPTECs were suspended in REBM media at a final density of 0.5 × 10^6^ cells/mL and 200 µL of the cell suspension was added to each well of the transwell plate. The monolayers were maintained at 37°C in an atmosphere of 5% CO_2_ with saturated humidity for up to 14 days. During this culture period, the medium was replaced at least three times each week.

### Transport studies using chips

Apparent permeability (P_app_) was experimentally derived in the apical (top channel) to basolateral (bottom channel) and basolateral to apical directions (**Supplementary Figure 1**). Following 8 days of culture, culture medium was aspirated from the donor channel inlet compartment and replaced by a substrate solution (1 µM for all the substrates and 5 µM for TEA) in supplemented CSC basal medium or supplemented REBM medium, depending on the direction permeability being assessed. For the inhibitor experiments, the inhibitors were dosed in both the top and bottom compartments for 1 hour at a flow rate of 80 µL/hour prior to the addition of the substrate. The total organic solvent contribution to the incubation was 1% (v/v) to avoid any cytotoxicity. After aspiration of the inhibitor pre-treatment medium and replacement with the dosing medium, the flow rate was increased to 600 µL/hour for 10 minutes to flush the channels. All media was aspirated from the outlet / collection reservoirs and the flow was reduced to 80 µL/hour to initiate the transport study. The incubation was conducted at 37°C for 3 hours and terminated by the collection of the donor and receiver samples in the outlet reservoirs. The samples were analyzed by liquid scintillation counter (LSC) or liquid chromatography-mass spectrometry (LC-MS/MS) to determine the concentration of probe substrates, in comparison to the dosed.

Prior to commencing these studies, loss of compound to the system components due to absorption and adsorption was characterized (**Supplementary Figure 2**). None of the 5 compounds evaluated absorbed or adsorbed significantly into the chip material.

### Transport studies using transwell

P_app_ was determined in the apical to basolateral and basolateral to apical directions. Following 8 and 14 days of culture, culture medium was aspirated, followed by the addition of substrate solution (1 µM for all the substrates and 5 µM for TEA) in the corresponding donor compartments to initiate the transport study. The incubation was conducted at 37°C for 2 hours and terminated by the collection of the donor and receiver samples. The samples were analyzed by LSC or LC-MS/MS to determine the concentration of probe substrates.

### LC-MS/MS

A Sciex Triple Quad™ 5500 LC-MS/MS system in positive ion mode was used to analyze the concentrations of non-radiolabeled substrates (AB Sciex, Framingham, MA). Liquid chromatography was conducted using Shimadzu (Kyoto, Japan) system controller (CBM-20A), pumps (LC-30AD), autoinjector (SIL-30AC), Acquity^®^ UPLC BEH C18 column (2.1 × 50 mm; 1.7 µM) (Waters, Milford, MA). Compounds were eluted in a linear gradient mobile phase mixture consisting of 0.1% formic acid in water (mobile phase A) and 0.1% formic acid in acetonitrile (mobile phase B).

### vRNA-seq Analysis

Total RNA was isolated from cells in the Kidney-Chip (4 individual samples obtained on Day 14) and transwells (4 samples obtained on Day 14) using buffer RLT plus and RNeasy^®^ Mini Kit (Qiagen, Hilden, Germany) according to the manufacturer’s protocol. The extracted RNA was analyzed using the Illumina^®^ HiSeq^®^ 4000 platform with maximum read length 2×150 bp paired-end.

The sequencing depth was ∼50M paired-end reads/sample. The average quality score was >35 for all samples. The Trimmommatic v.0.36 was used to trim the sequence reads to remove the poor-quality sequences and nucleotides. The trimmed reads were mapped to the Homo sapiens reference genome GRCh38 (available on ENSEMBL) using the STAR (Spliced Transcripts Alignment to a Refrerence) aligned v.2.5.2b. The generated BAM files were used to calculate the unique gene hit counts using the feature counts from the Subread package v1.5.2 Notably; only unique reads that fell within the exon regions were counted.

With exclusion of those poorly expressed genes across all samples, DGE analysis was performed between the two groups (Kidney-Chip cells and transwell cells) using the “DESeq2” R package by Bioconductor. Widely accepted threshold (adjusted p-value <0.01 and |log2FoldChange| > 1.6) was applied to the DGE analysis (Love *et al*., 2014).

### KEGG pathway analysis and GO term enrichment analysis

The 1,839 up-regulated genes in Kidney-Chips were used to run KEGG pathway analysis and Gene Ontology (GO) enrichment analysis. Both analyses were performed using the popular Database for Annotation, Visualization and Integrated Discovery (DAVID) v6.8 (**https://david.ncifcrf.gov/home.jsp) (**Huang *et al*., 2009a; b) (Ashburner *et al*., 2000; Kanehisa and Goto, 2000).

### Immunocytochemistry

The cells in the chips were fixed with 4% paraformaldehyde in PBS and incubated for 30 minutes at room temperature, which was followed by incubation with a permeabilization buffer composed of 0.125% Triton X-100 in PBS for 10 minutes. The cells were then blocked with solution containing 10% goat serum and 2% BSA in PBS for 1 hour at room temperature. The primary antibodies were added to cells and incubated overnight at 4°C, followed by incubation with secondary antibodies for 1 hour at room temperature. The cells were counterstained with Hoechst-33324 (Thermo Fisher, Waltham, MA). The primary antibodies used in this study were AQP-1 (Abcam, ab15080), sodium potassium ATPase (Abcam, ab7671), and vascular endothelial (VE)-Cadherin (Thermo Fisher, BS-0878R).

### Determination of Permeability and Inhibition Potency

Apparent permeability (*P*_*app*_) was assessed using methods described previously for static culture systems (Tran *et al*., 2004; Heikkinen *et al*., 2010) with adaptations for a flow-based system:

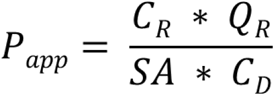

where *P*_*app*_ is the apparent permeability in units of cm/s, *SA* is the surface area of the co-culture channel (0.17cm^2^), Q_R_ is the fluid flow rate in the receiving channel in units of cm^3^/s, C_R_ is the concentration in the receiving channel in any consistent units, and C_D_ is the concentration in the dosing channel in any consistent units. As the recovery of all the compounds were greater than 90%, the loss of mass due to non-specific cell binding or material absorption was considered negligible (Error! Reference source not found.). Efflux ratios were then calculated by dividing the apparent permeability in the basal to apical direction by the apparent permeability in the apical to basal direction.

Inhibitor IC_50_ values were determined from victim compound efflux ratio data versus the corresponding inhibitor concentrations. A value of one was subtracted from all efflux ratios to express ratios in terms of transport above passive diffusion, which would be represented by an efflux ratio of one. These values were then normalized to the uninhibited efflux ratio for the victim-perpetrator combination, thereby expressing results as a percentage of the uninhibited active transport efflux ratio. This resulted in a dose-response curve with the uninhibited case represented by 100% transport and the completely inhibited case represented by 0%. The Hill Equation was then fit to the data to determine an IC_50_ for each perpetrator-inhibitor pairing at a victim dosing concentration of 1µM.

### Clearance Estimation

An estimation of renal clearance was made by following the classical equation (Russel *et al*., 2002; Lee and Kim, 2004; Feng *et al*., 2010; Morrissey *et al*., 2013; El-Kattan and Varma, 2018):

Essentially, the equation sums the predicted contributions to clearance of glomerular filtration and proximal tubule active transport but neglects any possible distal tubule reabsorption,

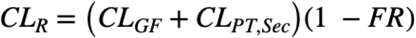

where *CL*_*R*_ denotes total renal clearance, *CL*_*GF*_ is clearance due to glomerular filtration alone, *CL*_*PT,Sec*_ denotes clearance due to proximal tubule secretion or active transport, and *FR* denotes the fraction of compound reabsorbed in the distal tubule. The exclusion of the fraction reabsorbed term results in an upper estimate of renal clearance for compounds where reabsorption is significant/appreciable. Reabsorption of xenobiotics in the distal tubule is driven by passive transport and tends toward an equilibrium between the unionized concentration in the filtrate and the unionized, non-protein bound fraction in the blood (Goodman *et al*., 2011). Fraction reabsorbed was estimated assuming achievement of this equilibrium, based on physiologically relevant blood and glomerular filtrate acidity (pH) ranges, whether the compound were acids or bases, the acid dissociation constant (pKa) of the compound, and filtrate and blood flow rates (**Supplementary Figure 4**). Unless otherwise indicated, distal tubule reabsorption was insignificant and ignored. Clearance due to glomerular filtration was calculated as:

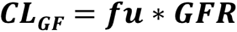

where *fu* denotes the fraction of compound unbound to plasma proteins and GFR is the Glomerular Filtration Rate (∼120 mL/min in healthy young adults per 1.73 m^2^ of body surface area). Clearance due to proximal tubule secretion was estimated using the general form:

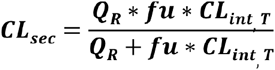

where Q_R_ is the renal perfusion rate (∼1200 mL/min in healthy adults per 1.73 m^2^ body surface area) and CL_int,T_ is the intrinsic clearance due to proximal tubule transport. Intrinsic transport clearance was estimated as the fractional increase in compound concentration of the glomerular filtrate within the proximal tubule compared to initial glomerular filtrate concentration entering the proximal tubule, relative to the concentration of unbound compound in systemic circulation. Assuming equilibrium is achieved between active secretion transport and passive diffusion back into the blood stream, intrinsic transport clearance takes the form:

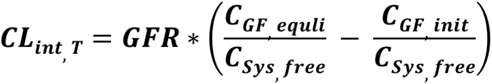

where *C*_*GF,equil*_ denotes the concentration of the molecule of interest in the proximal tubule glomerular filtrate at equilibrium, *C*_*Sys,free*_ denotes the concentration of free or unbound compound in the blood in systemic circulation, C_GF,init_ denotes the initial glomerular filtrate concentration or the concentration in the proximal tubule immediately following glomerular filtration *(see* **Supplementary Figure 4** *for full information*). Assuming the initial concentration of compound in the filtrate (C_GF,init_) is at equilibrium with the concentration of free compound in systemic circulation (*C*_*Sys,free*_) *in vivo* and substituting in the *in vitro* derived efflux for the ratio of the equilibrium concentration of compound in the filtrate (*C*_*GF,equil*_) versus the concentration of free compound in systemic circulation (C_GF,equil_/C_Sys,free_), the equation simplifies to the form:

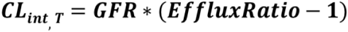

See supplemental method for full derivation. Substrate/victim clearance values were calculated both in the presence and absence of inhibitor/perpetrator and compared directly with literature-reported values for total/systemic human clearance.

### Modeling Clinical Data

A single-compartment clearance model was used to simulate the pharmacokinetics of all clinical studies analyzed and predict clinical study outcomes. Specifically, the model was used to predict shifts in clearance and area under the plasma concentration curve for the given dosing regimen of a victim-perpetrator versus dosing with only the victim/substrate. Model results were then compared to the reported clinical study results. Clinical data was pulled from the University of Washington Drug Interaction Database, with information regarding dosing regimen (dose mass, dosing interval, dosing duration) used as inputs to the predictive model. Other model inputs from *in vivo* data included either inhibitor concentration at steady-state (*C*_*ss*_) or inhibitor peak concentration (*C*_*max*_), which were taken from the database, depending on whether inhibitor was administered as a series of doses or a single dose. The model also took as inputs the volume of distribution of the victim compound (*V*_*d,ss*_) and the fraction unbound to serum proteins (*fu*) for both victim and perpetrator, which were pulled from two databases (Goodman *et al*., 2011; Lombardo *et al*., 2018). Finally, the relationship between perpetrator concentration and victim efflux ratio was ascertained from the Kidney-Chip results, which directly corresponded to a change in victim clearance. Model predictions were compared to reported clinical outcomes from these studies.

### Statistical analysis

Experiments were performed in triplicate for each sample per group. All error bars represent standard deviations of the mean, with errors propagated following standard practice. The statistical significance was calculated using student’s t-test and a p-value of <0.05 was considered to be statistically significant.

## Authorship contributions

Participated in research design: Nookala, Luo, McKenzie, Hamilton, Jang

Conducted experiments: Nookala, He, Ronxhi, Jeanty, Jadalannagari, Park, Jang

Contributed new reagents or analytic tools:

Performed data analysis: Nookala, Ronxhi, Sliz, Manatakis, Jang

Wrote or contributed to the writing of the manuscript: Nookala, Sliz, Manatakis, Lavarias, Luo, Ronxhi, Park, Jang

