## Supplemental figures for "Assessment of Human Renal Transporter Based Drug-Drug Interactions Using Proximal Tubule Kidney-Chip"

### Supplementary Material

**Supplementary Figure legends**

**Supplementary figure 1:** Schematic of the transport of solute molecules when dosed in the top channel and the bottom channel independently.

**Supplementary figure 2:** Compound recovery in Kidney-chip: (A) Adefovir, digoxin, metformin, PAH, and TEA were added to the apical and basal sides independently. Recovery is calculated by measuring the compound concentrations from the outlets and the inlets. Percentage of the compound recovered when dosed on the apical side and the basal side are represented by black solid bar and white solid bar, respectively (n=3 independent chips).

**Supplementary figure 3: Comparison of the transporter activities on kidney-chip versus transwell after 8 and 14 days in culture:** (A) The apparent permeability from the apical (A) to basal (B) direction (P_app_(A-B)) and permeability from the basal to apical direction (P_app_(B-A)) and the efflux ratio (P_app_(A-B)/P_app_(B-A)) were determined for various probe substrates, including digoxin (P-gp), metformin (MATEs and OCT2), tetraethyl ammonium (OCT2), and p-amino hippuric acid (OAT1) on day 8 and 14 using kidney-chip and transwell. Apparent permeability from A to B and B to A directions are represented in gray and black solid bars, respectively. The efflux ratios on day 8 and 14 are presented in blue solid bars (n = 3 independent chips).

**Supplementary figure 4: Derivation of renal clearance**

**Supplementary Figure 1**

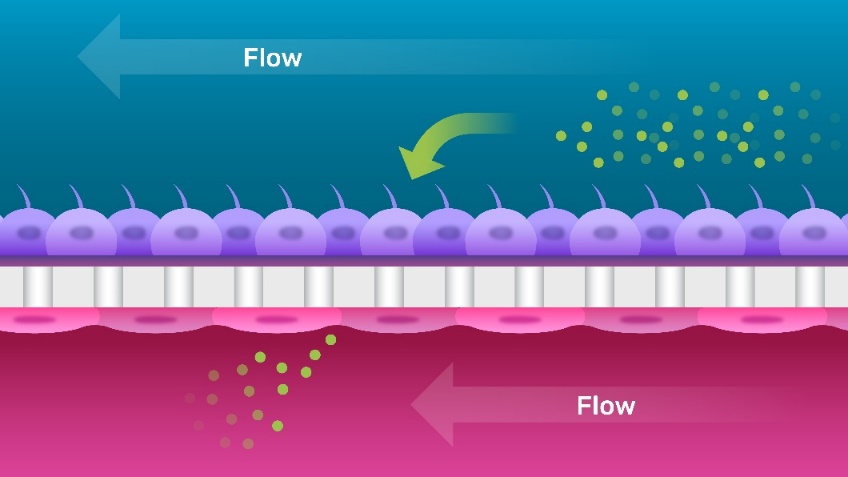

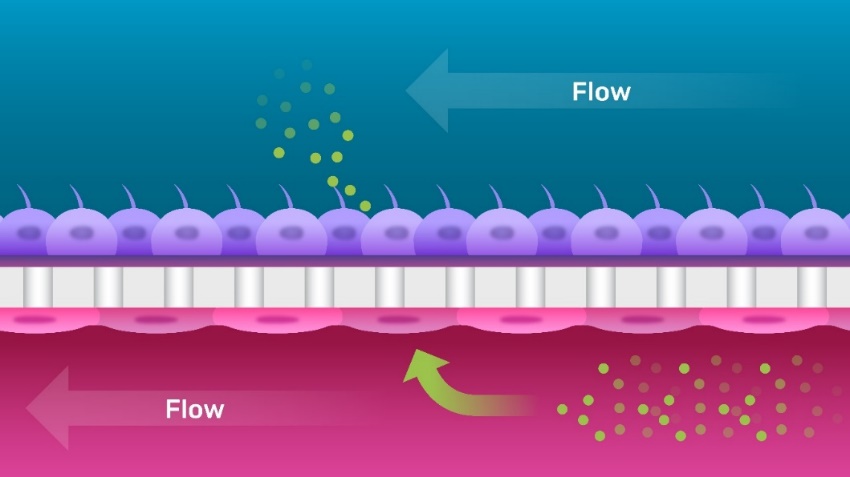

Top Channel Dosed

Bottom Channel Dosed

**Supplementary Figure 2**

Compound Recovery

Percent of Compound Recovered

|  | Apical Dosing |
| --- | --- |
| Adefovir | 95.942 |
| Digoxin | 92.6851 |
| Metformin | 95.609 |
| PAH | 98.004 |
| TEA | 94.1617 |

**Supplementary Figure 3**

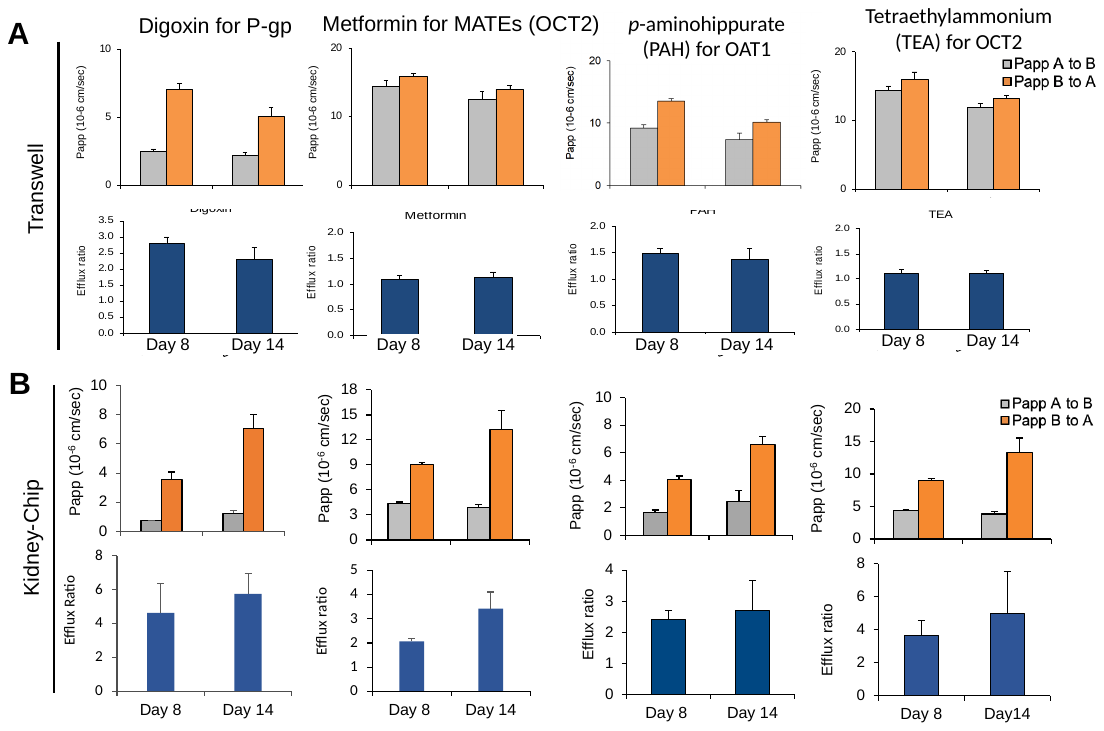

**Supplementary Figure 4**

Derivation of Whole Kidney Clearance from *in vitro* Kidney Proximal-Tubule-Chip

1. **Systemic Clearance**
   a. Definition of systemic clearance:

$${CL}_{sys,b}=\frac{Rate of Elimination}{{Concentration}_{b}}$$

where *CL_sys,b_* is the total human blood clearance (in mL/min or mL/min/kg of body weight), *Rate of Elimination* is the change in mass of the molecule in the body with time (in mg/min or mg/min/kg), and *Concentration_b_* is the concentration of the molecule in the blood (in mg/mL).

b. Systemic clearance as a summation of organ clearances:

$${CL}_{sys,b}={CL}_{R}+{CL}_{H}+{CL}_{Other}$$

where *CL_sys,b_* is the total human blood clearance, *CL_R_* is total renal clearance, *CL_H_* is total hepatic clearance, and *CL_Other_* is the summation of clearance from all other routes, which for many xenobiotics will be negligable.

c. Whole blood vs plasma clearance:

$${CL}_{sys,b}=\frac{{CL}_{sys,p}}{\left( BP \right)}$$

Where CL­_sys,p_ is total human plasma clearance (in mL/min) and BP is the ratio between compound concentration contained within whole blood and concentration in the plasma fraction – plasma concentration being the biological fluid most often sampled in a clinical setting.

2. **Total Renal Clearance**
   a. Governing equation for renal clearance:

$${CL}_{R}=\left( {CL}_{GF}+{CL}_{Sec} \right)\left( 1 -FR \right)$$

where *CL_R_* is total renal clearance also known as kidney organ-level clearance, *CL­_GF_* is clearance due to glomerular filtration alone, *CL_Sec_* is the clearance due to proximal tubule active transport also known as secretory clearance, and *FR* is the fraction of compound reabsorbed in the distal tubule.

b. For the purposes of this analysis, we examine the two cases that bound maximum and minimum kidney clearance. First, if there is minimal reabsorption, as is the case for compounds with low passive permeability, as may be identified by a low octanol-water partition coefficient (add Varma reference), then the fraction reabsorbed can be assumed to be nearly zero and the equation for whole kidney clearance simplifies to only the contributions of glomerular filtration and proximal tubule secretion. In the second case, where permeability of the compound of interest is relatively high, compound will tend toward an equilibrium in the distal tubule between the unionized concentration in the urine and the unionized, unbound concentration in the renal blood flow. In reality, this equilibrium often is never achieved due to slow passive transport, and the observed distal tubule passive reabsorption may in fact be negligible. Nevertheless, this equilibrium state can be determined using only physicochemical parameters of the compound and represents the bounding case in which estimates of total kidney clearance would be a minimum. Coupled with upper estimates of whole kidney clearance from the glomerulus and proximal tubule, this lower estimate is of great utility in defining the range of clearances that may be observed for a given compound in a clinical setting.
c. In the case where equilibrium is achieved within the proximal tubule between the filtrate and the kidney blood supply and where the compound is not significantly reabsorbed in the distal tubules, resulting in the fraction reabsorbed approaching zero, the equation for renal clearance simplifies to:

$${CL}_{R,max}=\left( {CL}_{GF}+{CL}_{Sec,equil} \right)$$

where CL_R,max_ is the maximum renal clearance and represents the upper bound for renal clearance for a given compound and CL_Sec,equil_ is the contribution to clearance from proximal tubule secretion, if equilibrium has been achieved between active and passive transport processes.

d. Within the distal tubule, compound could be entirely or partially reabsorbed via passive transport. The extent of this reabsorption is dependent on both the lipophilicity and ionization state of the molecule. In the limiting case, equilibrium is achieved between the unionized fraction of the compound within the tubule and the unionized, unbound fraction within the kidney blood flow. This results in an estimate of the minimum whole kidney clearance, as determined exclusively by distal tubule reabsorption:

$${CL}_{R,min}={CL}_{DT,equil}$$

where *CL_DT,equil_* is the minimum clearance after distal tubule reabsorption and represents the lower bound of renal clearance for a given compound.

3. **Estimation of Glomerular Clearance**
   a. Governing equation for glomerular clearance:

$${CL}_{GF}=fu*GFR$$

where *CL_GF_* is the clearance from glomerular filtration, *fu* is the fraction of compound unbound to plasma proteins, and *GFR* is the Glomerular Filtration Rate (~120mL/min in healthy young adults per 1.73m^2^ of body surface area).

b. Glomerular clearance represents the baseline clearance of the kidney for xenobiotics, with proximal tubule clearance increasing the concentration of the compound in the filtrate or urine via active transport, and distal tubule reabsorption decreasing the concentration for the compound in the filtrate via passive transport.

4. **Estimation of (Maximum) Proximal Tubule Clearance**
   a. Clearance due to proximal tubule clearance takes the general form:

$${CL}_{Sec}=\frac{Q_{R}*fu*{CL}_{int,T}}{Q_{R}+fu{*CL}_{int, T}}$$

where *Q_R_* is the renal perfusion rate (~1200 mL/min in healthy adults per 1.73m^2^ body surface area) and *CL_int,T_* is the intrinsic clearance due to transport - that is to say, the parameter characterizing the maximum rate of secretory clearance in the kidney, without the limitations of protein binding or kidney perfusion.

b. As filtrate flows through the proximal tubule, active transport from the blood increases the concentration of the compound of interest in the glomerular filtrate/urine. The maximum intrinsic ability of the proximal tubule to clear compound in this manner is set by the equilibrium condition, where the rate of active (and passive) transport into the filtrate is equal to the rate of passive transport back into the blood stream. In the non-kidney-perfusion-limited case, the maximum possible clearance from the proximal tubule is simply a function of the fractional increase in filtrate compound concentration over the initial concentration, which is set by glomerular filtration. The maximum fractional increase is caused when active transport is at equilibrium with passive transport:

$${CL}_{int,T}=GFR*\left( \frac{C_{GF,equil}}{C_{Sys,free}}- \frac{C_{GF,init}}{C_{Sys,free}} \right)$$

where *C_GF,equil_* is the concentration of the molecule of interest in the proximal tubule glomerular filtrate at equilibrium, *C_Sys,free_* is the concentration of free or unbound compound in the blood in systemic circulation, C_GF,init_ is the initial glomerular filtrate concentration or the concentration in the proximal tubule immediately following glomerular filtration. It is assumed that the concentration of the molecule of interest in the glomerular filtrate, within the proximal tubule immediately following glomerular filtration, is initially the same as the free fraction in blood. That is:

$$C_{GF,int}=C_{Sys, free}$$

resulting in simplification of the equation to:

$${CL}_{int,T}=GFR*\left( \frac{C_{GF,equil}}{C_{Sys,free}}- 1 \right)$$

c. C_GF,equil_/C_Sys,free_ can be calculated via *in vitro* assays, and is equivalent to an efflux ratio (which is defined further below):

$${CL}_{int,T}=GFR*\left( Efflux Ratio-1 \right)$$

As can be seen an efflux ratio greater than 1.0 will result in an increase in clearance over the baseline clearance governed by glomerular filtration alone. An efflux ratio less than 1 is not physiologically relevant and would not be expected in an *in vitro* context as this would otherwise indicate active transport from the filtrate/urine into the blood stream.

d. The efflux ratio for a given compound across a tissue barrier may be assessed *in vitro* by measuring the directional apparent permeability of that compound. This is achieved by alternately dosing the apical or basal reservoir in the case of a transwell or the channel in the case of an Organ-Chip, quantifying transported molecule over time in both directions, calculating an apparent permeability based on the measured concentration, then dividing the apparent permeabilities by one another:
   i. Net flux of molecules across a membrane or tissue barrier defined as:

$${Flux}_{net}=Active Transport+Diffusion$$

Or

$${Flux}_{net}={Flux}_{AB,active}+{Flux}_{BA,active}+{Flux}_{D}$$

where Flux_net_ is the net flux of molecule across the membrane, Flux_AB,active_ is the flux from apical to basal reservoir due to active transport, Flux_BA,active_ is the flux from basal to apical reservoir due to active transport, and Flux_D_ is the net flux across the membrane due to passive transport (or diffusion).

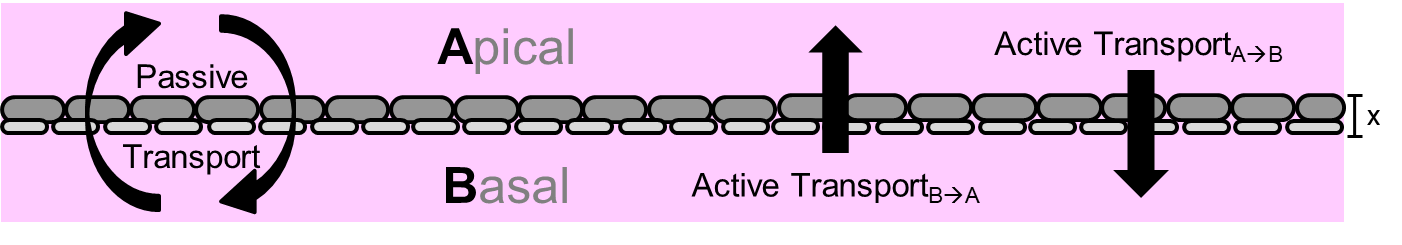

**Figure 1:** Transport across a biological membrane.

ii. Assuming Fickian Diffusion and first-order rates of active transport at steady-state the net flux is calculated as:

$${Flux}_{net}=K_{AB}\left[ A \right]-K_{BA}\left[ B \right]+D_{D}\frac{\left[ A \right]-\left[ B \right]}{\Delta x}$$

where K_AB_ is the active transport rate constant in the apical to basal direction, K_BA_ is the active transport rate constant in the basal to apical direction, [A] is the concentration of the molecule of interest in the apical reservoir, [B] is the concentration of the molecule of interest in the basal reservoir, D_D_ is the diffusivity of the molecule through the tissue layer, and Δx is the thickness of the tissue layer.

iii. Redefining the equation in terms of a P_app, passive_, or the passive permeability of the tissue:

$$P_{app}= \frac{D_{D}}{\Delta x}$$

yields:

$${Flux}_{net}=K_{AB}\left[ A \right]-K_{BA}\left[ B \right]+P_{app,passive}\left( \left[ A \right]-\left[ B \right] \right)$$

iv. Rearranging the equation yields:

$${Flux}_{net}=\left( K_{AB}{+ P}_{app,passive} \right)\left[ A \right]-\left( K_{BA}{+ P}_{app,passive} \right)\left[ B \right]$$

v. Defining the two direction-dependent apparent permeabilities:

$$\left( K_{AB}{+ P}_{app,passive} \right)= P_{app,AB} AND \left( K_{BA}{+ P}_{app,passive} \right)= P_{app,BA}$$

yields:

$${Flux}_{net}=P_{app,AB}\left[ A \right]-P_{app,BA}\left[ B \right]$$

vi. By experimentally dosing each reservoirs independently (apical or basal), which results in one reservoir experiencing a concentration of nearly “0” over the duration of the study, and assessing flux of the molecule of interest via concentration recovered in the non-dosed reservoir, each P_app,x_ can be assessed individually. For example, if the concentration in the basal compartment is set to “0”, P_app,AB_ can be directly assessed as a function of the flux across the membrane and the dosed concentration in the apical compartment as follows:

$$\left[ B \right]=0$$

then

$${Flux}_{net}=P_{app,AB}\left[ A \right]$$

and

$$P_{app,AB}= \frac{{Flux}_{net}}{\left[ A \right]}$$

vii. Efflux ratios are then simply computed as the apparent permeabilities for the two case where 1) the compound is dosed only in the apical chamber and 2) the compound is only dosed in the basal chamber:

$$Efflux Ratio= \frac{P_{app,BA}}{P_{app,AB}}$$

e. The efflux ratio also represents the concentration ratio between the two chambers at equilibrium:
   i. At equilibrium, the net flux across the membrane or tissue barrier “0”:

$${Flux}_{net}=P_{app,AB}\left[ A \right]-P_{app,BA}\left[ B \right]=0$$

and

$$P_{app,AB}\left[ A \right]=P_{app,BA}\left[ B \right]$$

ii. Rearranging and solving for the concentration ratio:

$$\frac{\left[ A \right]}{\left[ B \right]} = \frac{P_{app,BA}}{P_{app,AB}} = Efflux Ratio$$

which can be directly substituted into the equation for intrinsic transport clearance to obtain intrinsic transporter or proximal tubule clearance.

5. **Estimation of (Minimum) Clearance Due to Distal Tubule Reabsorption**
   a. Maximum clearance due to distal tubule reabsorption takes the same general form as proximal tubule secretion:

$${CL}_{DT,equil}=\frac{Q_{R}*fu*{CL}_{int,DT}}{Q_{R}+fu{*CL}_{int, DT}}$$

where *Q_R_* is the renal perfusion rate (1200 mL/min, in healthy adults) and *CL_int,DT_* is the maximum intrinsic clearance due to renal reabsorption in the distal tubule - that is to say, the parameter characterizing the limit of distal tubule reabsorption the kidney, without the limitations of protein binding or kidney perfusion.

b. Similarly, the intrinsic clearance within the distal tubule parallels the form of intrinsic transport clearance:

$${CL}_{int,DT}=GFR*\left( {Efflux Ratio}_{Effective,DT} \right)$$

where *Efflux Ratio_Effective,DT_* is the effective efflux ratio achieved at equilibrium across the distal tubule – blood interface, which can be assessed by *in silico* modeling. Unlike intrinsic transport clearance, the contribution of glomerular filtration and proximal tubule clearance do not need to be taken into account, since reabsorption attenuates clearance by these mechanisms, rather than adding to them and because here we consider the equilibrium case. Said another way, we assume that distal tubule reabsorption is complete and that equilibrium between renal blood and urine is not hindered or rate-limited by the diffusion across the tissue barrier.

c. Reabsorption of xenobiotics in the distal tubule is driven by passive transport and approaches equilibrium between the unionized concentration in the filtrate and the unionized, non-protein bound fraction in the blood. It is important to note again that this equilibrium may not be attained in the distal tubule. Indeed, important factors, like degree of lipophilicity (logP) will determine the propensity of compounds to pass freely through the distal tubule barrier and the speed at which they transit. Nonetheless, the equilibrium state does provide a prediction of the maximum distal tubule reabsorption possible, which is achieved at equilibrium:

$$\left[ Distal \right]_{unionized}= \left[ Blood \right]_{unionized, unbound}$$

where [Distal]_unionized_ is the concentration of unionized compound in the distal tubule (in the filtrate) and [Blood]_unionized,unbound_ is the concentration of unionized, unbound compound in the blood perfusing the kidneys. Protein binding in the filtrate is assumed to be negligible due to nearly complete filtration of albumin and alpha-1 acid glycoprotein in the glomerulus.

d. Percent ionization of a given molecule is defined by the whether the compound is an acid or base, the *pKa* of the compound, and *pH* of the biological fluid:

$$\left( \% unionized \right)= \left( 1 - \frac{1}{1+{10}^{x*\left( pH -pka \right)}} \right)$$

, where ${x=}_{+1 (base)}^{-1 (acid)}$

e. Applying this equation to the filtrate within the distal tubule (urine) and renal blood supply, respectively:

, where ${x=}_{+1 (base)}^{-1 (acid)}$

$$\left[ Distal \right]_{unionized}=\left[ Distal \right]*\left( 1 - \frac{1}{1+{10}^{x*\left( {pH}_{urine} -pka \right)}} \right)$$

and

$\left[ Blood \right]_{unionized, unbound}=fu*\left[ Blood \right]*\left( 1 - \frac{1}{1+{10}^{x*\left( {pH}_{blood} -pka \right)}} \right)$

, where ${x=}_{+1 (base)}^{-1 (acid)}$

f. Setting the unionized, unbound concentration in the blood equal to the unionized concentration in the urine and solving for the ratio between distal tubule concentration vs blood concentration yields an effective distal tubule efflux ratio:

$${Efflux Ratio}_{effective,DT} =\frac{\left[ Distal \right]}{\left[ Blood \right]}=f_{u}*\frac{\left( 1 - \frac{1}{1+{10}^{x*\left( {pH}_{blood} -pka \right)}} \right)}{\left( 1 - \frac{1}{1+{10}^{x*\left( {pH}_{urine} -pka \right)}} \right)}$$

, where ${x=}_{+1 (base)}^{-1 (acid)}$

g. However, since protein binding is accounted for in the equation for total distal tubule reabsorption, the fraction unbound term is dropped from this equation in the assessment of distal tubule clearance:

, where ${x=}_{+1 (base)}^{-1 (acid)}$

$${Efflux Ratio}_{effective,DT,int} =\frac{\left[ Distal \right]}{\left[ Blood \right]}=\frac{\left( 1 - \frac{1}{1+{10}^{x*\left( {pH}_{blood} -pka \right)}} \right)}{\left( 1 - \frac{1}{1+{10}^{x*\left( {pH}_{urine} -pka \right)}} \right)}$$

where Efflux Ratio_effective,DT,int_ is the intrinsic effective distal tubule efflux ratio at equilibrium.

h. This efflux ratio can be entered into the equation for intrinsic distal tubule clearance, which is by definition independent of protein binding in the blood. As can be seen, an effective distal tubule efflux ratio less than the proximal tubule active transport efflux ratio will result in net reabsorption and estimates of clearance using this method represent a lower bound for kidney clearance. Efflux ratios of exactly 1 indicate that glomerular filtration represents the lower bound. Efflux ratios greater than the proximal tubule efflux ration are not physiologically relevant, as they would indicate active transport mechanisms in the distal tubule.
